# Selective enrichment of A-to-I edited transcripts from cellular RNA using Endonuclease V

**DOI:** 10.1101/522029

**Authors:** Steve D. Knutson, Jennifer M. Heemstra

## Abstract

Immunoprecipitation enrichment has significantly improved the sensitivity and accuracy of detecting RNA modifications in the transcriptome. However, there are no existing methods for selectively isolating adenosine-to-inosine (A-to-I) edited RNAs. Here we show that *Escherichia coli* Endonuclease V (eEndoV), an inosine-cleaving enzyme, can be repurposed to bind and isolate A-to-I edited transcripts from cellular RNA through adjustment of cationic conditions. While Mg^2+^ is required for eEndoV catalysis, it has also been shown that similar levels of Ca^2+^ instead promote binding of inosine without cleavage. Leveraging these properties, we observe that Ca^2+^-supplemented eEndoV is highly specific for inosine in RNA and exhibits low nanomolar binding affinity. We then demonstrate EndoVIPER (<u>Endo</u>nuclease <u>V</u> <u>i</u>nosine precipitation <u>e</u>n<u>r</u>ichment) as a facile and robust method to isolate A-to-I edited transcripts from cellular RNA. We envision the use of this approach as a straightforward and cost-effective strategy to enrich edited RNAs and detect A-to-I sites with improved sensitivity and fidelity.

## Introduction

Adenosine-to-inosine (A-to-I) RNA editing is an abundant post-transcriptional modification found in metazoans. Catalyzed by adenosine deaminases acting on RNAs (ADARs), this reaction alters both the chemical structure and hydrogen bonding patterns of the nucleobase.^1^ Inosines preferentially base pair with cytidine, effectively recoding these sites as guanosine. A-to-I editing is ubiquitous across most RNA types, directly altering amino acid sequences of protein-coding mRNAs as well as modulating the target specificities and biogenesis of small-interfering RNAs (siRNAs) and microRNAs (miRNAs), in turn affecting global gene expression patterns and overall cellular behavior. A-to-I RNA editing is crucial for transcriptomic and proteomic diversity, and continues to be implicated in a variety of biological processes, including embryogenesis, stem cell differentiation, and innate cellular immunity.^2–4^ Dysfunctional A-to-I editing has also been linked with numerous disease progressions, including autoimmune disorders, neurodegenerative pathologies, and several types of cancer.^5–6^

Sensitive and accurate identification of A-to-I sites is vital to understanding these broader biological roles, relationships with disease, and regulation dynamics. Contemporary mapping methods typically utilize high-throughput next-generation RNA sequencing (RNA-seq). Because inosine is decoded as guanosine by polymerases, raw cDNA readouts can be matched to a reference genome to detect A-G transitions as inosine sites.^7^ This approach has enabled a wide survey of A-to-I locations in a variety of different species and tissues, and yielded substantial insights into the overall editing landscape.^8^ However, this technique also requires significant investment of time and materials, and is further limited in both accuracy and sensitivity. In the absence of large amounts of matched high-quality RNA and DNA, it can be difficult to discriminate between true A-to-I editing sites and sequencing mis-calls, transcriptional errors, or single nucleotide polymorphisms (SNPs), requiring further *in vitro* assays to confirm or refute editing status. However, these downstream approaches are often only applicable to cultured cells, and can further introduce unintended off-target cellular changes. Moreover, despite the overall high number of A-to-I editing sites across the transcriptome, with millions of loci currently catalogued, inosine content is relatively low in cellular RNA, requiring large quantities of material and a significant number of RNA-seq reads to achieve sufficient sequencing depth and transcriptome coverage. This is further complicated by the observation that editing rates at individual sites can be highly variable or conditionally active, differing significantly across cell and tissue types, developmental states, and disease progression stages.^8–10^ Additionally, many key edited RNAs are only present in low abundance, yielding very few actual RNA-seq reads. In these cases, identification of A-to-I sites is possible, but actual editing rates cannot be quantified, as acquiring a statistically significant number of reads would require impractically large amounts of RNA or excessively high numbers of RNA-seq reads. Overall, while RNA-seq allows agnostic transcriptome-wide analysis, it is not ideal for probing A-to-I editing, as the vast majority of the data are filtered out and discarded, and thus a significant amount of throughput, depth, and coverage is wasted on unwanted RNA populations. Together, the present limitations in accurately and robustly characterizing A-to-I sites and RNA editing activity restricts our overall understanding of epitranscriptomic dynamics and regulation.

Enriching A-to-I edited transcripts from total RNA prior to analysis would largely overcome these challenges by depleting unedited RNAs that otherwise lead to “wasted” sequencing reads. Similar approaches for pulldown of modified bases has markedly improved the throughput and reliability in detecting several other epitranscriptomic and epigenetic modifications. Of note, immunoprecipitation (IP) of RNA and DNA using antibodies for *N*^6^-methyladenosine and 5-methylcytosine, respectively, has significantly reduced sample complexity prior to analysis, in turn drastically improving detection fidelity, sensitivity, and overall throughput.^11–12^ While a previous report detailed the generation of inosine-targeting polyclonal antibodies for enriching modified tRNAs, these were also found to cross-react with several other nucleobases, and this method has not been reproduced.^13^ We previously explored chemical labeling and enrichment of inosine using an acrylamide derivative, and while we demonstrated feasibility in modification and capture of inosine-containing RNAs, this method also displayed off-target reactivity with pseudouridine and uridine, limiting the enrichment efficiency.^14^ Currently, there are no robust methods for the affinity purification of inosine-containing transcripts. Herein we report EndoVIPER (<u>Endo</u>nuclease <u>V</u> <u>i</u>nosine <u>p</u>recipitation <u>e</u>n<u>r</u>ichment) as a novel method to selectively enrich A-to-I edited transcripts. EndoVIPER leverages the observation that supplementation of the enzyme with Ca^2+^ promotes binding rather than cleavage of A-to-I transcripts, enabling their isolation. We employ this technique in a magnetic immunoprecipitation workflow, validating EndoV specificity and efficiency in binding inosine in RNA. We then demonstrate EndoVIPER in enriching an A-to-I edited coding transcript from human brain cellular RNA, highlighting the utility of this approach to study A-to-I RNA editing in biological contexts.

## Results

### Endonuclease V specifically recognizes inosine in RNA and exhibits robust binding in the presence of Ca^2+^

Due to the previously reported difficulties in developing inosine-targeting antibodies, we instead searched for naturally-occurring proteins capable of recognizing and binding to inosine. We identified EndoV, a highly conserved nucleic acid repair enzyme found in all domains of life. In prokaryotes, EndoV is mainly responsible for detection of inosine resulting from oxidative damage in DNA, and cleaves after these deamination lesions to promote base excision repair.^15^ In humans and other metazoans, EndoV has been implicated in the metabolism of A-to-I edited RNAs.^16–17^ Thus, we hypothesized that if the cleavage activity could be selectively suppressed without compromising recognition and binding, then EndoV could be leveraged for enriching A-to-I edited RNAs. While human EndoV (hEndoV) appears to be a good candidate toward this goal, recent studies also identified variable substrate preferences and possible affinity toward both unedited double-stranded RNA (dsRNA) and ribosomal RNA (rRNA), properties which could be problematic for use in total RNA samples.^18^ Interestingly, these reports also showed that *Escherichia coli* EndoV (eEndoV) was both specific and highly active toward inosine in single-stranded RNA (ssRNA) and exhibited minimal substrate or sequence bias.^16–17^ These observations, as well as the commercial availability of a purified recombinant enzyme, encouraged us to explore eEndoV for the pulldown and enrichment of A-to-I edited transcripts.

Structural analyses of several orthologs have revealed that EndoV requires Mg^2+^ as a cofactor for inosine recognition and catalysis of strand scission (Fig. S1).^19^ Similar studies have also found that replacing Mg^2+^ with Ca^2+^ facilitates binding of EndoV to inosine substrates, yet does not support catalysis.^20^ While the exact mechanism underlying this observation is unknown, it is likely that differences in electronics and coordination chemistry between the two metals are key factors. In any case, we hypothesized that supplementing eEndoV with Ca^2+^ would enable enrichment of inosine-containing RNAs (Fig. 1a). To test this, we synthesized a pair of Cy5-labeled oligoribonucleotides having either A or I in a defined position, and evaluated eEndoV activity in the presence of both cations. Consistent with previous reports, we observed not only specific cleavage activity towards inosine in ssRNA (RNA I) when benchmarked against a non-edited control (RNA A), but also an obligate Mg^2+^ requirement for cleavage (Fig. S2). We next sought to evaluate the effect of Ca^2+^ supplementation on the ability of eEndoV to bind and isolate inosine-containing ssRNA. Conveniently, the recombinant enzyme is genetically fused to a maltose-binding protein (MBP) tag, enabling us to design a magnetic IP workflow using anti-MBP functionalized beads, which we term EndoVIPER (<u>Endo</u>nuclease <u>V</u> <u>i</u>nosine precipitation <u>e</u>n<u>r</u>ichment, Fig. 1b). Using this method, we attempted pull down of both RNA A and RNA I in the presence of variable amounts of Ca^2+^, monitoring the initial, unbound (flowthrough) and elution fractions after washing (Fig. 1c, S3). Not surprisingly, omitting Ca^2+^ produced little binding of either oligonucleotide, corroborating the notion that Mg^2+^ aids in both recognition and cleavage of inosine substrates. Increasing amounts of Ca^2+^ from 0-10 mM improved binding efficiency substantially, approaching ~80% recovery with excellent selectivity (~350-fold over pulldown of RNA A). Interestingly, additional supplementation beyond 10 mM Ca^2+^ quickly decreased pulldown efficiency and selectivity (Fig. 1c-d, S2). While unconfirmed, these results may arise from electrostatic shielding of the negative charge of the phosphodiester backbone, preventing key amino acid residues from interacting with the nucleic acid substrate. Regardless, we identified 5 mM Ca^2+^ as ideal for maximizing recovery and selectivity, and moved forward with measuring the binding affinity of eEndoV for both RNA substrates using microscale thermophoresis (MST). Consistent with our IP results, we observed low nanomolar affinity for RNA I and no measurable binding to the off-target control (Fig 1e, S3).

**Figure 1.**
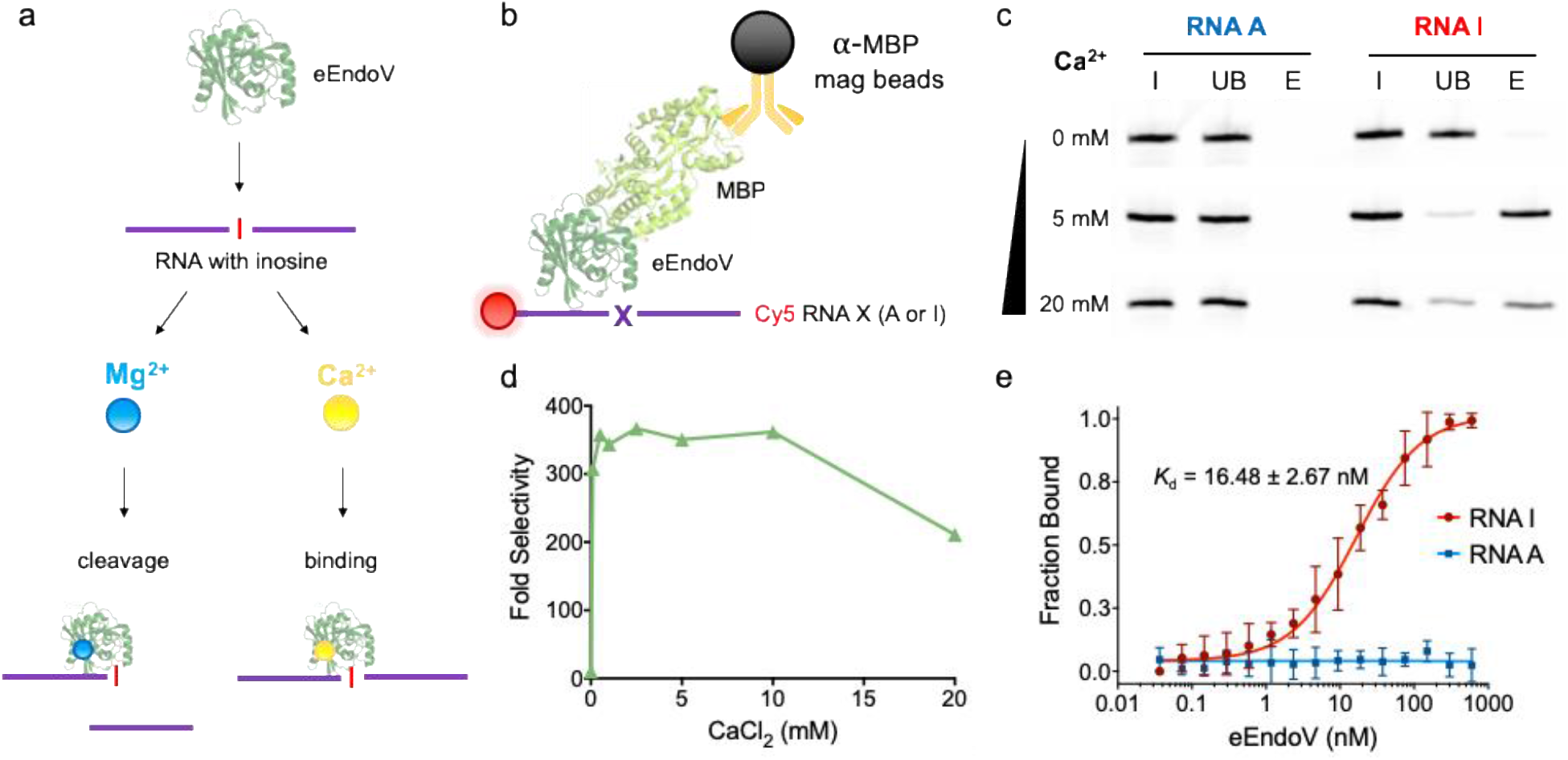
eEndoV binds inosine in RNA with high affinity and enables pulldown. a) Mg^2+^ or Ca^2+^ supplementation modulates eEndoV activity towards inosine-containing RNA substrates. b) Schematic of EndoVIPER with a Cy5 labeled RNA using recombinant eEndoV-MBP fusion protein and anti-MBP magnetic beads. c) Representative PAGE analysis of initial (I), unbound (UB) and eluate (E) EndoVIPER fractions, illustrating the effects of Ca^2+^ supplementation on pulldown efficiency. d) Identification of optimal Ca^2+^ concentrations by comparison of pulldown efficiency for A- and I-containing RNA. e) Quantification of eEndoV binding affinity towards RNA I (red) and RNA A (blue) using MST. Values and *K_d_* represent mean with 95% CI. (n = 3).

### EndoVIPER enables enrichment of A-to-I edited transcripts from cellular RNA

Encouraged by these results, we challenged our EndoVIPER protocol to enrich a naturally edited transcript from cellular RNA (Fig. 2a). To assess enrichment efficiency, we chose to track the *GRIA2* mRNA, an ionotropic glutamate receptor transcript that is highly edited in the brain.^9^ (Fig. 2b). We designed primers flanking the Q607R A-to-I recoding site and used quantitative polymerase chain reaction (qPCR) to measure enrichment of the amplicon. We observed ~45-fold enrichment of *GRIA2* from human brain mRNA compared to a mock magnetic bead control. However, when we applied this method in brain total RNA, we observed a nearly 10-fold drop in *GRIA2* enrichment efficiency (Fig 2c). We speculate this could be due to significantly higher content of other A-to-I edited transcripts that were quenching our pulldown. In particular, ~7-8 human tRNAs contain inosine at the wobble N34 position. Additionally, A-to-I sites have been identified in ribosomal RNA (rRNA).^21^ In either case, edited tRNA and rRNA would constitute a large molar proportion of our total RNA sample and potentially saturate the eEndoV binding sites.^22^ Regardless, we still observe statistically significant enrichment (~5-fold) of *GRIA2* from total RNA, demonstrating the method’s compatibility and efficacy in complex RNA samples. Together, these experiments show that EndoVIPER is capable of selectively isolating inosine-containing transcripts with high efficiency.

**Figure 2.**
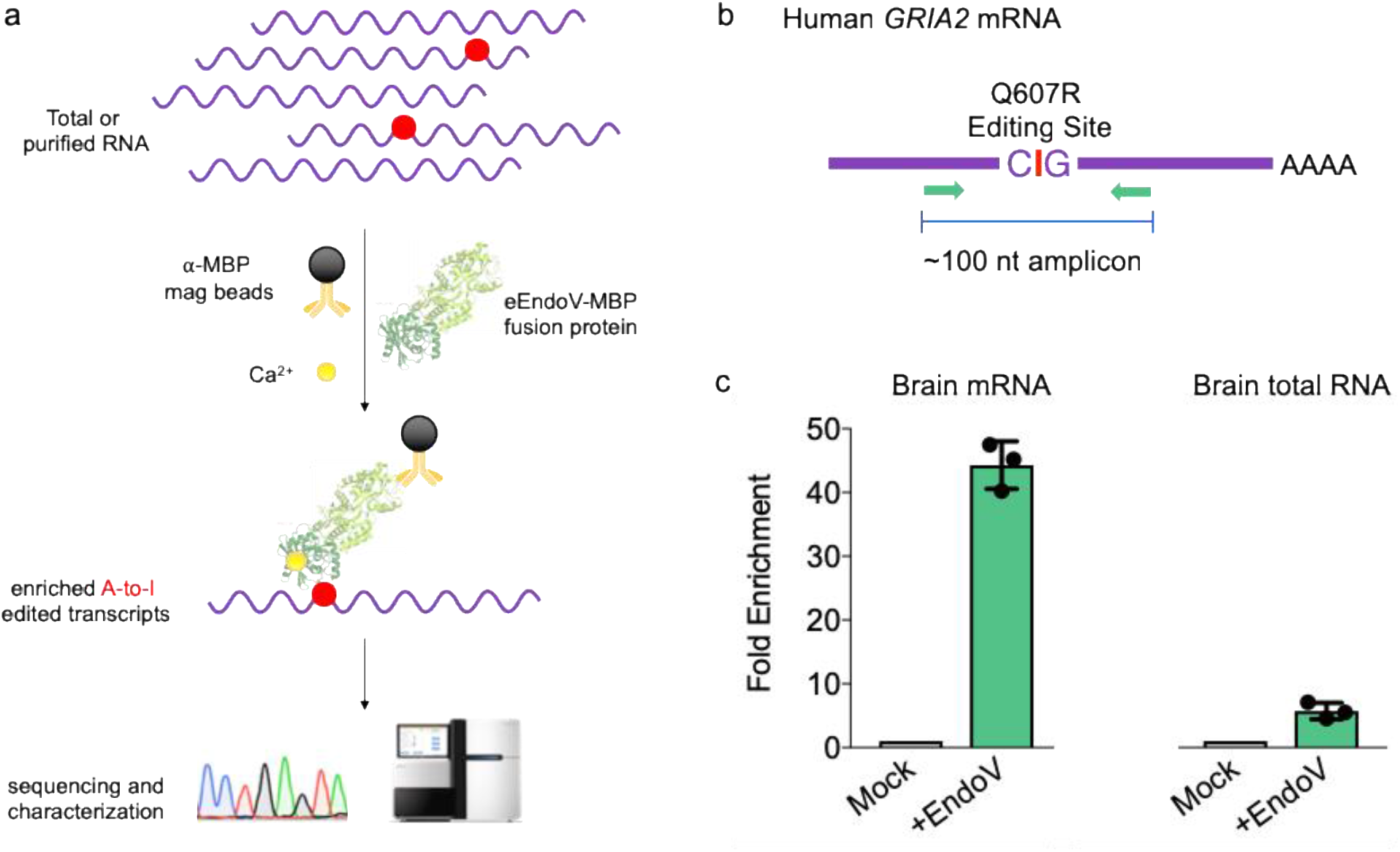
EndoVIPER enrichment of A-to-I edited transcripts from cellular RNA. a) EndoVIPER workflow for the enrichment and analysis of A-to-I edited RNAs b) *GRIA2* mRNA is a naturally edited transcript containing an A-to-I recoding site, allowing detection c) Fold-enrichment of GRIA2 using endoVIPER from brain mRNA and total RNA.

## Discussion

Herein we present EndoVIPER as a new method for the affinity pulldown of inosine-containing transcripts from RNA, overcoming existing limitations in the detection and characterization of A-to-I RNA editing in complex biological samples. Our approach is robust, displaying high affinity and selectivity for inosine in RNA. In addition, EndoVIPER exclusively utilizes low-cost, commercially available reagents with little to no modification. Further, this method is compatible with both purified and total cellular RNA, and fits well with existing library preparation protocols for both lower throughput analyses as well as next-generation RNA-seq.

While this study establishes a novel workflow for A-to-I enrichment, there remains opportunity for further improving EndoVIPER efficiency and selectivity. As described previously, EndoV is present across nearly all domains of life, offering a vast array of potential affinity scaffolds for use in subsequent iterations of this method. In addition, this approach could be further elaborated by in-depth protein engineering and directed evolution strategies to increase binding affinity and tune EndoVIPER selectivity for specialized sequence contexts or sample types. Further, a wide variety of purification tags and bioconjugation chemistries are available for affinity enrichment of biomolecules, offering additional avenues for optimization of this workflow. We anticipate that the overall ease of use and accessibility of this method will find utility in a number of empirical contexts, significantly improving our understanding of the dynamics and regulation of A-to-I RNA editing in a range of biological settings, in turn elucidating its role in normal cellular processes and its relationship with disease pathology.

## Methods

### RNA Oligoribonucleotides

All oligonucleotides used in this study were custom designed and purchased from Integrated DNA Technologies. Edited and non-edited controls were synthesized with a Cyanine5 (Cy5) label at the 5’ terminus as shown below.

RNA I 5’ Cy5 AAGCAGCAGGCU<u>**I**</u>UGUUAGAACAAU 3’

RNA A 5’ Cy5 AAGCAGCAGGCU<u>**A**</u>UGUUAGAACAAU 3’

### RNA Cleavage Assays

10 pmol of either RNA I or RNA A was incubated in the presence or absence of both Mg^2+^ at a 10 mM final concentration and/or 9pmol recombinant eEndoV (New England Biolabs) in a total volume of 10 μL. Final buffer conditions in all reactions were 10 mM Tris, 125 mM NaCl, 15 μM EDTA, 150 μM DTT, 0.025% Triton X-100, 30 μg/ml BSA, 7% glycerol, pH 7.4. Reactions were incubated for 1 hour at 25 °C, followed by a 10 min heat inactivation at 85°C. Reaction products were separated using 10% denaturing PAGE, and gels were imaged with a GE Amersham Typhoon RGB scanner using 635 nm excitation laser and the Cy5 670BP30 emission filter.

### EndoVIPER Magnetic IP Assays

For each binding test, 10 pmol of either RNA I or RNA A was combined with 30 pmol of eEndoV and variable amounts of CaCl_2_ (0, 0.1, 0.5, 1, 2.5, 5, 10 and 20 mM) in a total volume of 50 μL. Final buffer conditions were 10 mM Tris, 125 mM NaCl, 15 μM EDTA, 150 μM DTT, 0.025% Triton X-100, 30 μg/ml BSA, 7% glycerol, pH 7.4. Reactions were incubated at 25 °C for 30 min, after which a 3 μL sample (initial, I) was taken and set aside for later analysis. Separately, 70 μL of anti-MBP magnetic bead slurry (New England Biolabs) was washed extensively with a buffer containing 10 mM Tris, 125 mM NaCl, 7% glycerol, and variable amounts of CaCl_2_ (0, 0.1, 0.5, 1, 2.5, 5, 10 and 20 mM), pH 7.4. After washing, beads were resuspended in eEndoV-RNA samples and incubated at 25 °C for two hours with end-over-end rotation. Magnetic field was applied to beads and a 3 μL sample (unbound, UB) of the supernatant was saved for later analysis. Beads were washed extensively with respective buffers containing variable amounts of Ca^2+^, and resuspended in 50 μL 10 mM Tris, 125 mM NaCl, 0.5 mM EDTA, 47.5% formamide 0. 01% SDS, pH 7.4 and heated to 95 °C for 10 min. Magnetic field was applied and a 3 μL final sample (eluate, E) of the supernatant was taken of each reaction. Collected fractions were analyzed using 10% denaturing PAGE, and gels were imaged with a GE Amersham Typhoon RGB scanner. Densitometric quantification of bands was performed using ImageJ software. % Bound is expressed as a band intensity ratio of unbound versus initial fractions. % Recovered was defined as the intensity ratio of eluate versus initial fractions. Fold-selectivity was calculated as the ratio of RNA I versus RNA A recovery percentages.

### Microscale Thermophoresis (MST)

For each binding test, varying amounts of eEndoV were combined with 6 fmol of either RNA A or RNA I in a final volume of 20 μL and allowed to incubate for 30 min at 25 °C. Final buffer conditions in all samples were 10 mM Tris, 125 mM NaCl, 5 mM CaCl_2_, 15 μM EDTA, 150 μM DTT, 0.025% Triton X-100, 30 μg/ml BSA, 7% glycerol, pH 7.4. After incubating, samples were loaded into NT.115 standard glass capillaries. MST experiments were performed using a Nanotemper Monolith NT.115 Pico instrument. All measurements were analyzed using the Pico-RED filter with 12% LED intensity and 40% laser power. Data were fitted using GraphPad Prism 7 analysis software to determine *K*_d_ values. Binding tests were performed in triplicate in separate trials.

### qPCR measurement of EndoVIPER enrichment

1 μg brain mRNA (Takara Bio) or 10 μg of brain total RNA (Thermo Fisher Scientific) was briefly heated to 95 °C and incubated in the presence or absence (mock) of 30 pmol of eEndoV in a volume of 50 μL. Final buffer conditions were 10 mM Tris, 125 mM NaCl, 5 mM CaCl_2_, 15 μM EDTA, 150 μM DTT, 0.025% Triton X-100, 30 μg/ml BSA, 7% glycerol, pH 7.4. eEndoV-RNA reactions were allowed to incubate at 25 °C for 30 min. Separately, 70 μL of anti-MBP magnetic bead slurry (New England Biolabs) was washed extensively with a buffer containing 10 mM Tris, 125 mM NaCl, 5 mM CaCl_2_, and 7% glycerol, pH 7.4. After washing, beads were resuspended in eEndoV-RNA samples and incubated at 25 °C for two hours with end-over-end rotation. Beads were then washed extensively with 10 mM Tris, 125 mM NaCl, 5 mM CaCl_2_, and 7% glycerol, and resuspended in nuclease free water. Beads were heated to 95 °C for 20 min. Magnetic field was applied and the supernatant was collected, 0.2 μm filtered, and ethanol precipitated. The RNA pellets were then resuspended in 6 μL nuclease free water, mixed with 20 pmol of GRIA2 reverse primer (5’ CCACACACCTCCAACAATGCG 3’), heated to 70 °C for 10 min, and cooled on ice for 5 min. First strand cDNA synthesis was performed for one hour at 42 °C using M-MuLV Reverse Transcriptase (New England Biolabs). Reactions were then mixed with 10 pmol of GRIA2 forward primer (5’ GGGATTTTTAATAGTCTCTGGTTTTCCTTGGG 3’) and reverse primer and 1X iTaq Universal SYBR Green Supermix (BioRad). qPCR was monitored with a Roche Lightcycler 96 instrument, using the following PCR program: 94 °C for 3 min, followed by 45 cycles of (94 °C for 15 s, 60 °C for 30 s, 68 °C for 30 s), 68 °C for 5 min. Fold enrichment was determined from raw threshold cycle (Ct) values and defined as (2^-ΔΔCt^), where ΔΔCt = (Ct EndoVIPER) -(Ct mock). All mock and EndoVIPER pulldowns were performed independently in triplicate. PCR products were purified using the Monarch PCR and DNA cleanup kit (New England Biolabs), and electrophoresed on 1 % agarose gel to verify amplicon size.

