## Supplemental Figures for "Selective enrichment of A-to-I edited transcripts from cellular RNA using Endonuclease V"

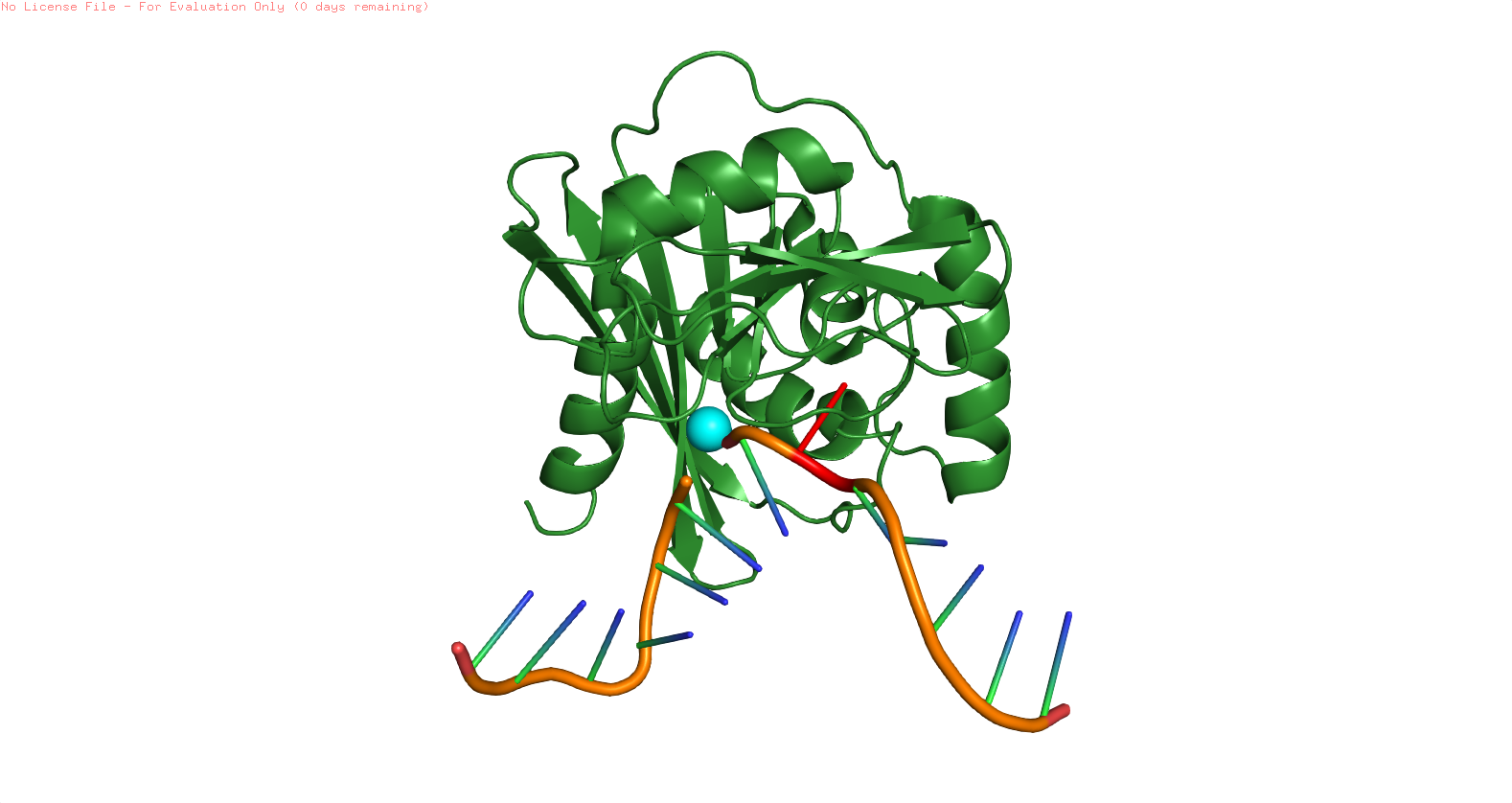

**Supplementary Figure 1.** a) Crystal structure (PDB 2W35) of eEndoV (green) complexed with ssDNA (orange), illustrating recognition of inosine (red) in a nucleic acid substrate. Mg^2+^ is shown in cyan adjacent to cleavage site.

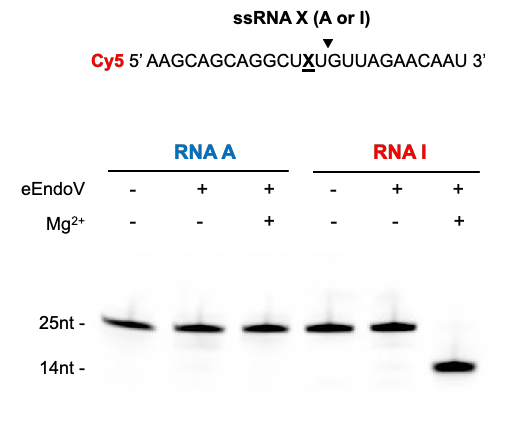

**Supplementary Figure 2.** Test oligoribonucleotide sequences with putative cleavage site (arrow) and PAGE analysis of reactions with eEndoV illustrating specificity toward RNA I and confirming Mg^2+^ requirement for cleavage.

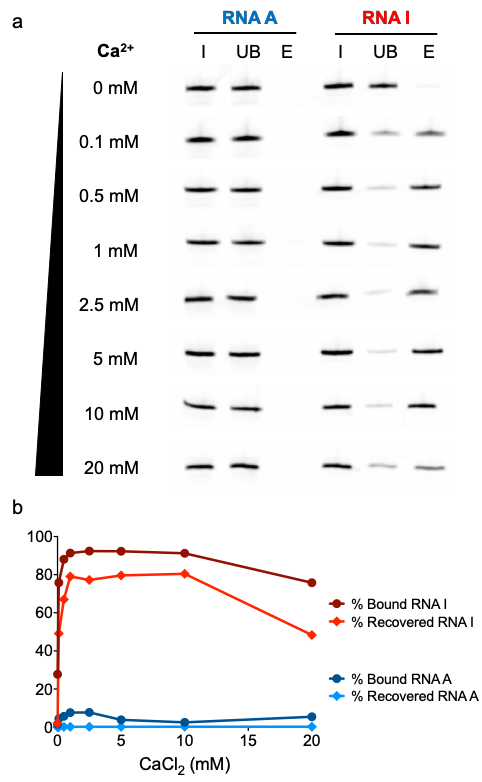

**Supplementary Figure 3.** a) Full immunoprecipitation and PAGE analysis of eEndoV enrichment using variable amounts of Ca^2+^. Initial (I), unbound (UB) flowthrough and eluate (E) fractions were analyzed with 10% denaturing PAGE. b) Densitometric analysis of PAGE results to estimate endoVIPER binding and recovery efficiencies in both RNA substrates as a function of Ca^2+^ concentration.

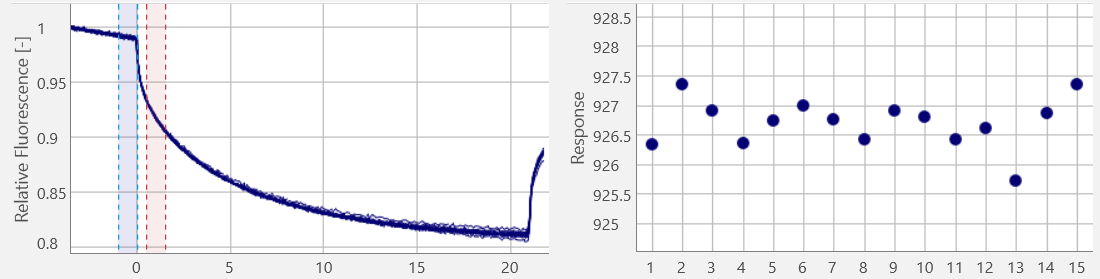

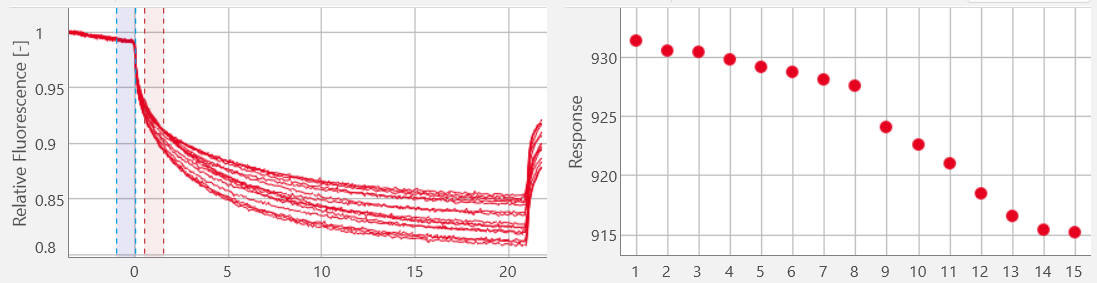

**Supplementary Figure 4.** Representative raw fluorescence and response traces using microscale thermophoresis (MST) to measure binding affinity between eEndoV and RNA A (blue) and RNA I (red).

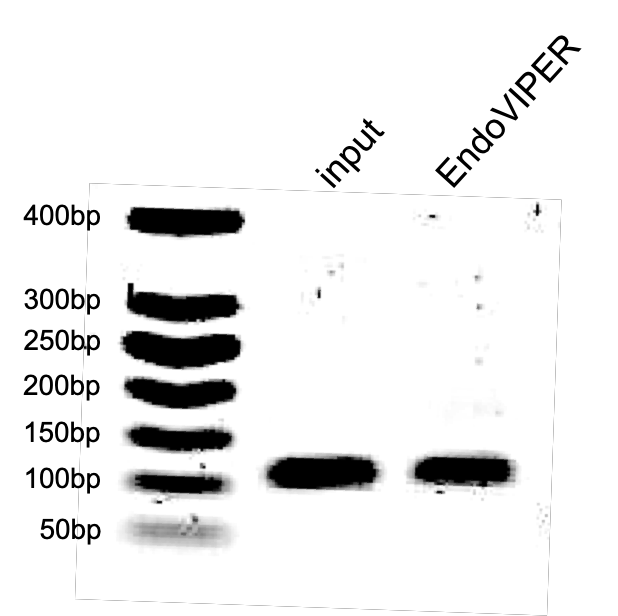

**Supplementary Figure 5.** Agarose gel analysis of GRIA2 PCR amplicons, verifying specific amplification in raw (input) and pulled down (EndoVIPER) RNA samples. Expected amplicon size is 103 nt.
